# Improved outcome and cost effectiveness in ablation of persistent atrial fibrillation under general anaesthetic

**DOI:** 10.1101/123448

**Authors:** Claire A Martin, James P Curtain, Parag R Gajendragadkar, David A Begley, Simon P Fynn, Andrew A Grace, Patrick M Heck, Kiran Salaunkey, Munmohan S Virdee, Sharad Agarwal

**Author notes:** **Corresponding author**: Claire A Martin, Papworth Hospital NHS Foundation Trust, Papworth Everard, Cambridge CB23 3RE, UK.

## Abstract

**Aims:** Outcome of persistent atrial fibrillation (AF) ablation remains suboptimal. Techniques employed to reduce arrhythmia recurrence rate are more likely to be embraced if cost-effectiveness can be demonstrated. A single-centre observational study assessed whether use of general anaesthesia (GA) in persistent AF ablation improved outcome and was cost-effective.

**Methods:** 292 patients undergoing first ablation procedures for persistent AF under conscious sedation or GA were followed. End points were freedom from listing for repeat ablation at 18 months and freedom from recurrence of atrial arrhythmia at one year.

**Results:** Freedom from atrial arrhythmia was higher in patients who underwent ablation under GA rather than sedation (63.9% vs 42.3%, HR 1.87, 95% CI: 1.23 to 2.86, p = 0.002). Significantly fewer GA patients were listed for repeat procedures (29.2% vs 42.7%, HR 1.62, 95% CI: 1.01 to 2.60, p = 0.044). Despite GA procedures costing slightly more, a saving of £177 can be made per patient in our centre for a maximum of 2 procedures if all persistent AF ablations are performed under GA.

**Conclusions:** In patients with persistent AF, it is both clinical and economically more effective to perform ablation under GA rather than sedation.

**What’s New?:**

- There is very little known regarding the clinical outcome of catheter ablation of AF under GA compared with sedation; to our knowledge there is one study only in paroxysmal AF and no studies examining cost effectiveness.
- This study shows that in patients with persistent AF, it is both clinical and economically more effective to perform ablation under GA rather than sedation.
- This study leads us to recommend the use of GA for the ablation of persistent AF. As PAF ablation is now increasingly being undertaken by single shot techniques which do not have the same requirements for analgesia and immobility, GA resources may be allocated for persistent AF ablation.

## Introduction

Atrial fibrillation (AF) is associated with increased mortality, congestive heart failure, and stroke. Catheter ablation (CA) has been established as a treatment option for patients with symptomatic, drug-refractory AF^1^. In paroxysmal AF (PAF) patients, the generation of circumferential lesions around veins to electrically isolate them from the rest of the LA is the cornerstone treatment^2^, which may currently be achieved by a range of ‘single-shot’ techniques as well as a ‘point-by-point’ technique.

Results of CA in patients with persistent AF are less satisfactory^3^. Pulmonary vein isolation (PVI) is still central; however, recurrent and chronic AF induces progressive electrical and tissue structural remodeling^4^. To interrupt this process, addition of a substrate based ablation approach may be employed, using a combination of linear lesions in the left atrium (LA) and focal ablation of complex fractionated atrial electrograms (CFAEs)^3,5^. For this reason, a ‘point-by-point’ approach which allows more tailored therapy, is more often employed for the CA of persistent AF. Whilst spontaneous reversion to sinus rhythm (SR) may be an endpoint of ablation, more commonly patients require electrical cardioversion at the end of the procedure.

CA may be performed under general anesthesia (GA) or sedation that may be conscious or deep. In the UK, an anaesthetist must be present to administer propofol for deep sedation and therefore this option is rarely chosen. The use of GA however, gives several potential benefits. There is greater comfort for the patient in a procedure that may last for several hours and involve some discomfort. They will therefore be less likely to move and distort the 3D electroanatomic geometry that has been created for the LA. The patients’ respiratory rate and depth are regulated, making respiratory movement of catheters inside the LA smaller and more predictable. The mapping system is also able to better calculate respiratory compensation. If sedation is administered or if the patient is in pain, their breathing is more likely to become erratic.

Anecdotally, operators find procedures under GA easier to undertake and feel that outcome is improved, but there is a lack of objective data supporting this. Cost is often cited by hospital management as a preventative factor in scheduling more GA lists. Therefore, this study sought to examine the outcome of patients undergoing persistent AF ablations under GA or conscious sedation, both in terms of arrhythmia recurrence and of the need for repeat ablations, and to calculate the cost effectiveness of both strategies.

## Materials and methods

### Study Design

This was an analysis of prospectively kept data of all cases of first catheter ablation for persistent AF ablation from the beginning of 2008 to the beginning of 2014 in a single large-volume centre. The study was approved by local ethics committee. Cases were excluded if they had had a previous ablation for either paroxysmal or persistent AF. All patients were over 18 years of age and all had symptomatic persistent AF (of over 7 days duration) refractory to at least one anti-arrhythmic agent. All patients were anticoagulated for at least 1 month pre-procedure and 3 months post-procedure, and given heparin to maintain an ACT of 300-350s during the procedure.

Patients underwent ablation either under conscious sedation with fentanyl and midazolam, or under GA. This was mainly related to the availability of GA provision on certain days of the week, and was not targeted to any particular subgroup of AF patients. In the GA group, after initiation of appropriate monitoring as prescribed by AAGBI standards, induction of anaesthesia was achieved by propofol 2-3mg/kg and remifentanil infusion 0.02-0.3mcg/kg/min. The airway was secured either by endotracheal intubation (facilitated by administering a non-depolarising muscle relaxant) or insertion of a laryngeal mask airway. Anaesthesia was maintained by target controlled infusion of propofol & remifentanil, guided by continuous depth of anaesthesia monitoring by the use of a BIS (Bi spectral index) monitor. In the conscious sedation group, intravenous doses of fentanyl and midazolam were administered by a nurse as directed by the physician to achieve moderate sedation.

AF ablation was performed by a point-by-point technique using Carto (Biosense Webster) or Ensite NavX (St Jude Medical) guidance. Contact force sensing was available only in cases performed at the end of the study period; when employed, a contact force of 5-40 g was targeted. Ablation strategy was pulmonary vein isolation (PVI) +/- additional substrate ablation in the form of roof or mitral isthmus line or CFAE ablation. The strategy employed was dependent on operator preference rather than case selection. The endpoint of ablation was PVI confirmed by abolition of PV potentials, effective linear ablation confirmed by bidirectional block across the roof of the LA and around the mitral valve isthmus, or CFAE ablation confirmed by elimination of all CFAE sites in the LA and coronary sinus. Atrial tachycardias were mapped and ablation attempted. If the patients were in AF or an atrial tachycardia at the end of the procedure, they were cardioverted.

Data was collected of the patient demographics, of the ablation strategy and of the clinical outcome. Clinical assessments, 12 lead ECGs and where appropriate 24 hr Holter monitors, were obtained at baseline and at 3, 6, 12 and 18 months after the ablation.

### Cost analysis

Equipment is listed as the price paid for it by the Trust and staff costs are calculated based on the salary per hour. Although between 2008 and 2014, two different mapping techniques were used, currently in our institution persistent AF ablation is almost entirely done using a point-by-point RF technique with Carto (Biosense Webster) mapping, and therefore this technique was costed for. Hotel services include the cost of the hospital bed, cleaning and food. Ward care includes the cost of the nursing and medical staff during the patient‘s inpatient stay. Costs are listed in Table 5 as a mean per patient; for example, only 7% of patients underwent post-procedure echocardiography, therefore the mean cost per patient of an echo is £2.94. Procedures under GA had the same ward based costs as those under sedation. They used all the equipment and staff as those under sedation plus additional anaesthetic equipment and staff, listed separately.

### Study outcomes

The primary study outcome was freedom from listing for repeat ablation procedures at 18 months. Due to the variable and often very long waiting lists for AF ablation during this time, this was felt to provide a better measure of the need for repeat ablation than freedom from the repeat ablation procedure itself. All patients put on the waiting list for an ablation did go on to have an ablation within a year of being put on the waiting list. Main secondary outcomes included freedom from any episodes of atrial arrhythmia at 1 year after one ablation procedure.

Atrial arrhythmia occurring within an initial 3 months blanking period after ablation was not counted, in accordance with the guidelines^6^. An episode of atrial arrhythmia was considered part of the outcome analyses if it lasted longer than 30 seconds and was documented by any form of monitoring, regardless of symptoms.

### Statistical Analysis

We estimated a sample size of 70 per arm would be needed to provide 80% power (p < 0.05, 1 sided) to detect a 50% change in the hazard ratio (HR) between groups, assuming a median freedom from atrial arrhythmia of 12 months and a follow-up of 18 months^7^.

Continuous variables are presented as mean ± standard deviation, and categorical data as counts or percentages. Analysis and comparisons of continuous data were performed using ANOVA, whilst the *χ^2^* test was used to compare categorical data. Survival was estimated using Kaplan Meier (KM) analyses. Cox proportional-hazards-models were used to explore univariate and multivariate predictors of events. Initial exploratory co-variates of age, gender, LA diameter, left ventricular ejection fraction (LVEF), presence of risk factors including hypertension, diabetes, stroke, vascular disease and congestive heart failure (CHF), CHA_2_DS_2_-VASc score, ablation technique, length of time in AF, and use of anti-arrhythmic drug post procedure were used. Multivariate models included terms with p-value of <0.1 at univariate analysis. A two-sided probability level of <0.05 was considered significant. All calculations were performed using SPSS 20.0 (IBM Software, USA).

## Results

### Patients

A total of 339 patients were identified who had first ablations for persistent AF between 2008 and 2013. Of these 47 were excluded from the study because of: a lack of information regarding their procedure or follow-up (19), the procedure being abandoned before any ablation took place (5), the patients not in fact having the procedure they were labelled as having (6), the patients having had a prior PAF ablation (17).

Out of the 292 included patients, 220 had procedures under sedation and 72 under GA. There was no significant difference in the numbers undergoing procedures under GA in each year in the time period studied. Mean doses of sedation given were 3.2 +/- 2.3 mg midazolam and 75 +/- 35 mcg fentanyl. Baseline characteristics were not significantly different between groups (Table 1). A followup period of up to 18 months was examined. All patients had clinical assessment and 12 lead ECGs performed at each clinic visit. If there was no documentation of atrial arrhythmia but the patient described any symptoms of palpitations, SOB or fatigue which might be attributable to a recurrence of AF or to an organised atrial arrhythmia, they underwent Holter monitoring to achieve ECG documentation of a symptomatic episode (occurred in 22% of total patients).

**Table 1.** Characteristics of patients included in the study. Plus–minus values are means ±SD. * denotes significant difference between groups (see main test)

|  |  | Sedation | GA | p-value |
| --- | --- | --- | --- | --- |
| **Characteristic** |  |  |  |  |
|  | Number | 220 | 72 |  |
|  | Age - yr | 59 +/- 9 | 60 +/- 10 | 0.463 |
|  | Male sex - no (%) | 181 (82) | 57 (79) | 0.588 |
|  | Time in persistent AF - months | 27 +/- 26 | 32 +/- 29 | 0.204 |
|  | Procedure duration - min | 179 | 184 | 0.349 |
|  | Ablation time - s | 2623 +/- 1323 | 2876 +/- 1335 | 0.778 |
|  | Nights in hospital | 1.1 +/- 0.7 | 1.0 +/- 0.8 | 0.331 |
| **Medical history - no (%)** | Hypertension | 77 (35) | 24 (33) | 0.884 |
|  | Diabetes | 1 (1) | 8 (9) | 0.286 |
|  | Vascular disease | 8 (4) | 3 (4) | 0.720 |
|  | Stroke/TIA | 17 (8) | 2 (2) | 0.261 |
|  | Heart failure | 16 (7) | 7 (10) | 0.438 |
| **CHA<sub>2</sub>DS<sub>2</sub>-VASc score - n (%)** | 0 | 90 (41) | 33 (46) | 0.245 |
|  | 1 | 76 (35) | 17 (24) |  |
|  | 2 | 32 (15) | 14 (19) |  |
|  | >2 | 22 (10) | 8 (11) |  |
| **Baseline medications - n (%)** | Beta-blocker | 89 (41) | 29 (40) | 0.670 |
|  | Ca channel blocker | 18 (8) | 3 (4) | 0.426 |
|  | Flecainide | 51 (23) | 17 (24) | 0.742 |
|  | Amiodarone | 35 (16) | 9 (13) | 0.846 |
|  | Dronedaron | 3 (1) | 1 (1) | 1.00 |
|  | Sotalol | 29 (13) | 4 (6) | 0.128 |
| **LA diameter - n (%)** | normal | 108 (49) | 42 (58) | 0.773 |
|  | mildly dilated | 62 (28) | 18 (25) |  |
|  | moderately dilated | 36 (16) | 9 (13) |  |
|  | severely dilated | 14 (6) | 3 (4) |  |
| **EF - n (%)** | normal | 161 (73) | 54 (75) | 0.997 |
|  | mildly impaired | 28 (13) | 9 (13) |  |
|  | moderately impaired | 19 (9) | 6 (8) |  |
|  | severely impaired | 11 (5) | 3 (4) |  |
| **Ablation technique** | PVI | 67 (30) | 18 (25) | 0.280 |
|  | PVI + lines | 63 (29) | 22 (31) |  |
|  | PVI + CFAEs | 11 (5) | 4 (6) |  |
|  | PVI + lines + CFAEs | 79 (36) | 28 (39) |  |

### Outcomes

At 12 months, freedom from atrial arrhythmia lasting longer than 30s after one ablation procedure, with or without the use of antiarrhythmic medications, was present in 42.3% of patients who underwent ablation under sedation and 63.9% who underwent isolation under GA (Table 2). There was a significant difference in freedom from atrial arrhythmia (HR 1.87, 95% CI: 1.23 to 2.86, p =0.002) (Fig 1a). There was no difference in procedure time and ablation time between groups.

**Table 2.** Outcome as a function of sedation or GA use

|  | GA (n = 72) | CS (n = 220) | p-value |
| --- | --- | --- | --- |
| **Freedom from listing for repeat ablation procedure at 18 months (%)** | 51 (70.8) | 126 (57.3) | 0.044 |
| **Freedom from any atrial arrhythmia at 1 yr (%)** | 46 (63.9) | 93 (42.3) | 0.002 |

**Figure 1.**
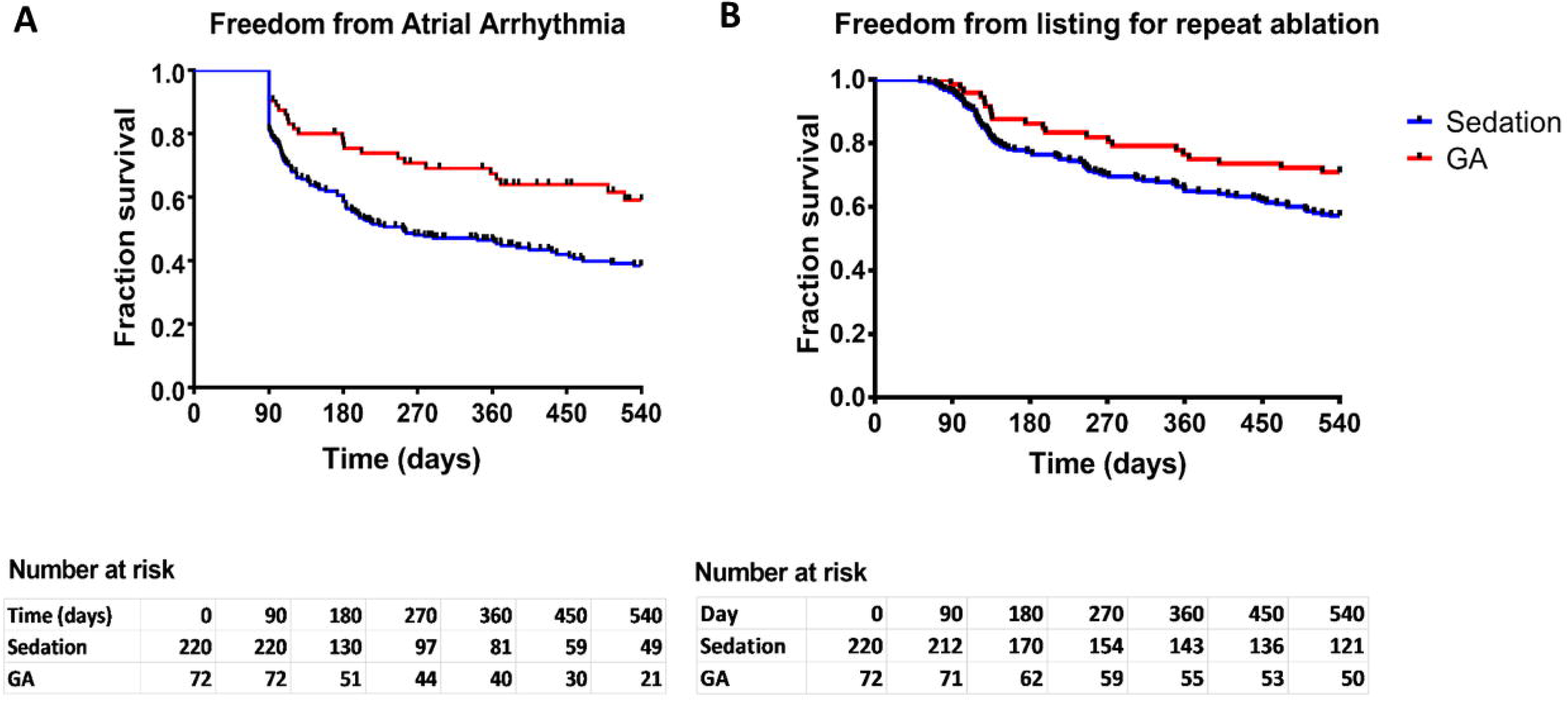
Kaplan Meier survival curves depicting (A) freedom from atrial arrhythmia recurrence after one ablation procedure and (B) freedom from listing for repeat ablation.

During the 18 month follow up period, 42.7% of patients undergoing AF ablation under sedation were listed for a repeat ablation procedure, compared with 29.2% under GA (Table 2). There was a significant difference in freedom from listing for a repeat ablation between groups (HR 1.62, 95% CI: 1.01 to 2.60, p = 0.044) (Fig 1b).

Of the patients who had recurrence of arrhythmia but did not undergo repeat ablation, 39% did not because they had only very occasional recurrences of PAF, 20% were either having frequent episodes of PAF or were still in persistent AF but had felt subjectively better, and in 17% it was felt that there was a low chance of success from further procedures. Other reasons included the patient declining a further procedure, the patient not having felt any better when in sinus rhythm, the patient having a DCCV and then maintaining SR, the patient having an AV node ablation or a MAZE procedure at the same time as a CABG, or having died of other causes before being put on the waiting list (Fig 2).

**Figure 2.**
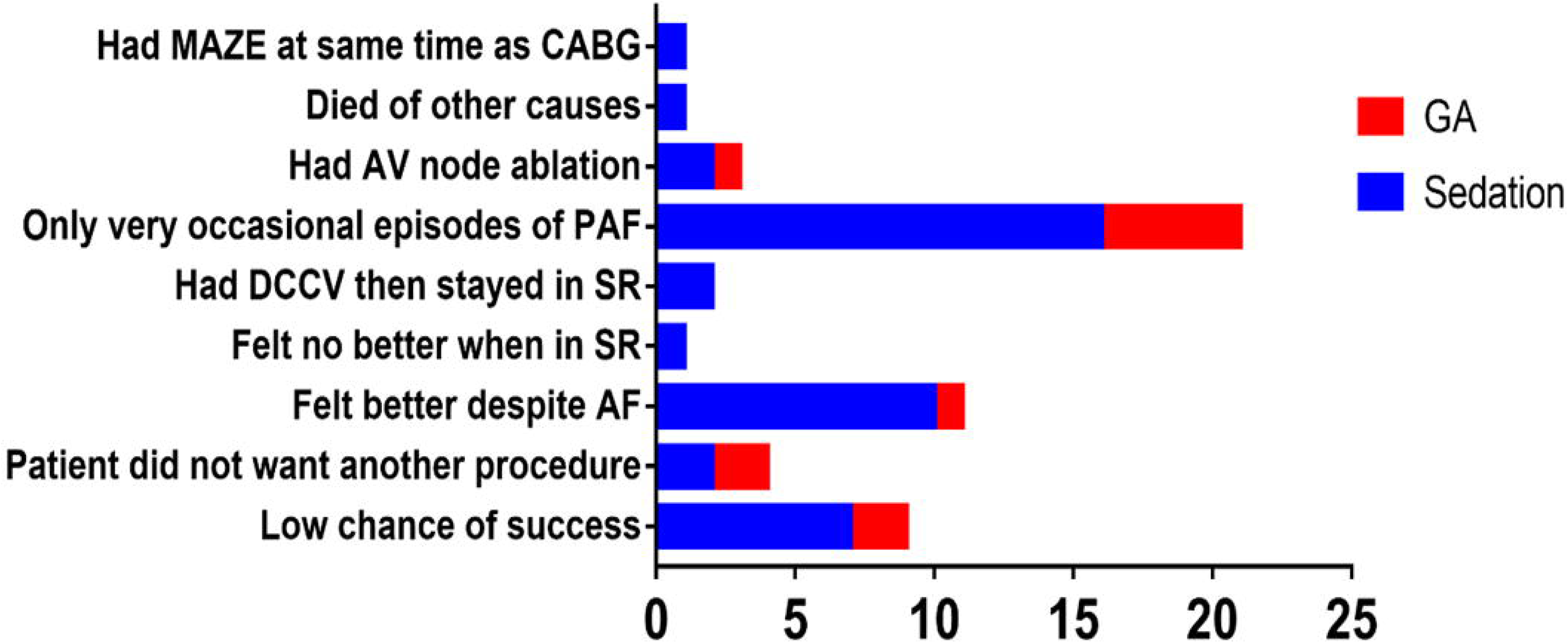
Reasons patients with AF recurrence were not added to waiting list for repeat ablation

Freedom from any atrial arrhythmia and freedom from listing for repeat ablation was analysed stratified by age, gender, ablation technique, use of GA, presence of risk factors including stroke, diabetes, CHF, hypertension and PVD, CHA_2_DS_2_-VASc score, LA diameter, EF, time in AF, and whether an anti-arrhythmic (amiodarone, dronedarone, flecainide or sotolol) was used post procedure.

Univariate Cox regression analysis found the following factors to increase likelihood of freedom from atrial arrhythmia at the p < 0.1 level (Table 3): decreasing age, normal or only mildly dilated LA, decreasing time in AF pre-procedure, use of GA and use of an antiarrhythmic drug post procedure. In multivariate analysis, there was higher freedom from atrial arrhythmia with use of GA over sedation, as well as for decreasing age, normal or only mildly dilated LA and decreasing time in AF preprocedure.

**Table 3.** Factors influencing freedom from atrial arrhythmia at 12 months. Bold text denotes significant difference between groups (see main text)

| Predictors | Univariate |  |  | Multivariate |  |  |
| --- | --- | --- | --- | --- | --- | --- |
|  | HR | 95% CI | p-value | HR | 95% CI | p-value |
| Age /year | 0.983 | 0.996 - 1.000 | **0.051** | 0.960 | 0.96 - 0.99 | **0.012** |
| Gender - male | 0.797 | 0.542 - 1.173 | 0.250 |  |  |  |
| Normal/mildly dilated LA | 1.460 | 1.017 - 2.083 | **0.033** | 1.510 | 1.020 – 2.273 | **0.041** |
| Normal LV function | 1.170 | 0.851 – 1.608 | 0.334 |  |  |  |
| Hypertension | 1.054 | 0.683 - 1.32 | 0.756 |  |  |  |
| Diabetes | 0.884 | 0.577 - 2.218 | 0.721 |  |  |  |
| Stroke | 0.706 | 0.663 - 3.024 | 0.366 |  |  |  |
| Vascular disease | 0.604 | 0.540 - 3.704 | 0.386 |  |  |  |
| CHF | 1.302 | 0.451 - 1.31 | 0.333 |  |  |  |
| CHA2DS2-VASc score $\geq 2$ | 0.807 | 0.559 – 1.166 | 0.254 | | | |
| Use of GA | 1.721 | 0.378 – 0.892 | **0.013** | 1.600 | 1.01 - 2.55 | **0.048** |
| Length of time in AF/months | 0.995 | 0.999 - 1.011 | **0.086** | 0.995 | 0.990 - 1.000 | **0.041** |
| Anti-arrhythmic post-procedure | 1.039 | 0.994 – 1.086 | **0.091** | 1.287 | 0.814 – 2.033 | 0.280 |
| PVI only vs PVI + substrate | 0.775 | 0.549 – 1.094 | 0.147 |  |  |  |

In univariate analysis, decreasing age and use of GA over conscious sedation increased the likelihood of freedom from listing for repeat ablation at the p < 0.1 level (Table 4). In multivariate analysis, the only factor significant at the p <0.05 level was decreasing age, but use of GA over sedation was significant at the p < 0.1 level.

**Table 4.** Factors influencing freedom from listing for repeat ablation at 18 months. Bold text denotes significant difference between groups (see main text)

| Predictors | Univariate |  |  | Multivariate |  |  |
| --- | --- | --- | --- | --- | --- | --- |
|  | HR | 95% CI | p-value | HR | 95% CI | p-value |
| Age /year | 0.972 | 0.954 – 0.990 | **0.003** | 0.972 | 0.954 – 0.991 | **0.003** |
| Gender - male | 0.992 | 0.623 - 1.579 | 0.974 |  |  |  |
| Normal/mildly dilated LA | 0.778 | 0.508 – 1.192 | 0.248 |  |  |  |
| Normal LV function | 0.942 | 0.653 – 1.360 | 0.750 |  |  |  |
| Hypertension | 1.154 | 0.781 - 1.703 | 0.472 |  |  |  |
| Diabetes | 1.054 | 0.490 - 2.263 | 0.894 |  |  |  |
| Stroke | 1.114 | 0.518 – 2.393 | 0.782 |  |  |  |
| Vascular disease | 1.657 | 0.526 – 5.216 | 0.388 |  |  |  |
| CHF | 1.017 | 0.515 – 2.010 | 0.960 |  |  |  |
| CHA2DS2-VASc score $\geq 2$ | 1.438 | 0.728 – 2.839 | 0.296 | | | |
| Use of GA | 1.634 | 1.018 – 2.623 | **0.042** | 1.504 | 0.924 – 2.446 | **0.099** |
| Length of time in AF/months | 1.003 | 0.996 - 1.010 | 0.434 |  |  |  |
| Anti-arrhythmic post-procedure | 1.057 | 0.731 – 1.529 | 0.769 |  |  |  |
| PVI only vs PVI + substrate | 0.889 | 0.595 – 1.328 | 0.567 |  |  |  |

### Adverse events

2 strokes, 3 tamponades and 1 groin haematoma occurred in patients undergoing ablation under sedation. 1 groin haematoma and 1 reactive pericardial effusion occurred in patients undergoing ablation under GA. There were a further 5 cases under sedation that were hindered by airway problems, patients becoming agitated or with uncontrolled pain.

### Financial implications

Having established that outcomes in terms of freedom of atrial arrhythmia and repeat ablations are significantly better with procedures under GA rather than sedation, the cost implications of performing all persistent AF ablations under GA was calculated (Table 5). GA procedures were slightly more expensive at £4406.68 compared to £4115.15 for procedures under sedation. However, taking into account the higher redo rates for procedures under sedation, the total cost per patient after a maximum of two procedures is £178 less with procedures under GA (Table 6). This may be an underestimate of the true saving, as some patients go on to have multiple procedures.

**Table 5.**
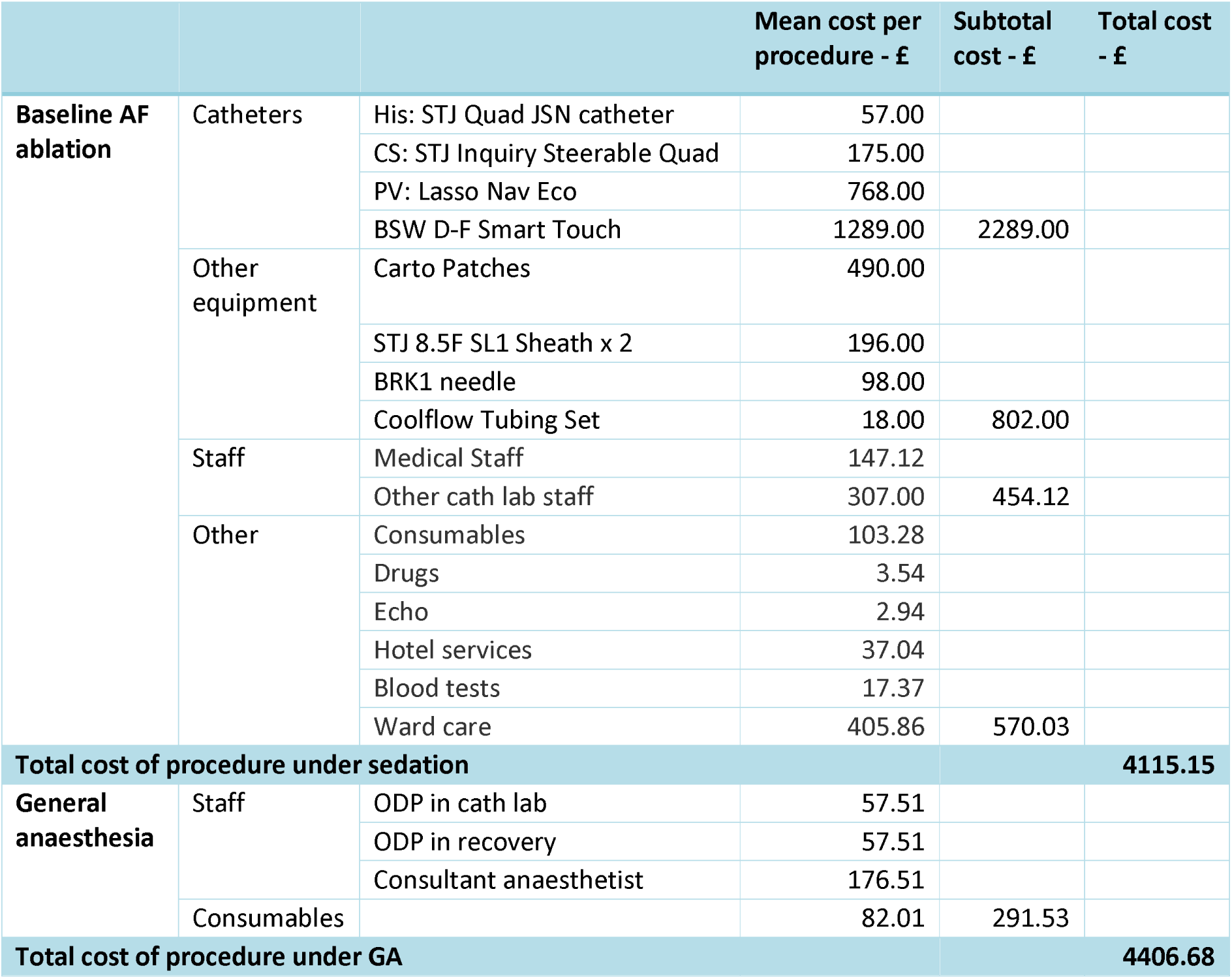
Cost of AF ablation using point-by-point technique with Carto mapping, with and without GA

**Table 6.**
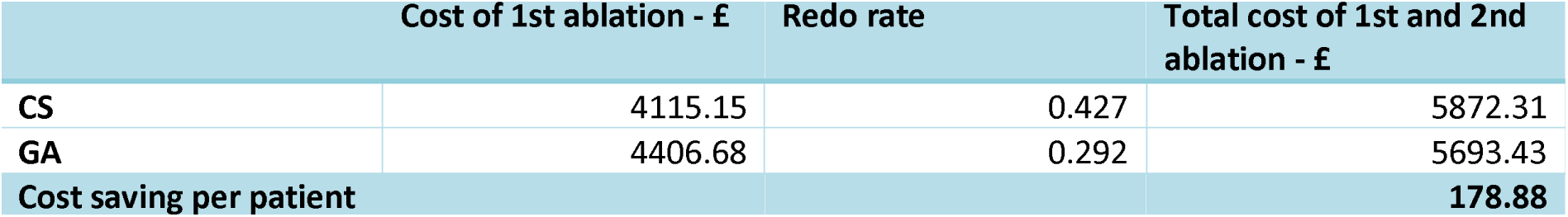
Cost saving per patient by performing ablations for persistent AF under GA rather than sedation

## Discussion

This is an observational study of persistent AF ablation in a single large-volume centre. We found that outcome of ablation is improved if the procedure had been conducted under GA rather than conscious sedation. There is little objective evidence for this yet published in the literature. Di Biase et al^8^ have demonstrated better outcomes for PAF ablation after a single procedure, with a reduction in the prevalence of PV reconnection observed at the time of repeat ablation and also reduced fluoroscopy and procedure times. A retrospective analysis of PAF ablation^9^ has also showed improved outcome after GA over sedation. However, currently first time PAF ablations are usually limited to PVI alone, with procedures often undertaken by a ‘single shot’ rather than ‘point-by-point’ technique. This does not require the patient to remain as still to maintain the accuracy of the 3D electro-anatomical mapping, and the procedure is also shorter and less painful. One could argue that the use of GA is of particular use in persistent AF ablation, where more extensive substrate ablation means that ‘point-by-point’ ablation is usually employed, procedures last longer, and where a DCCV is often required near the end of the procedure after the main ablation has been completed but before vein isolation can be verified.

Kumar et al. recently showed the benefit of ablating during apnoea (possible in patients under GA), comparing contact-force parameters during RF pulses between apnoea and ventilation^10^. During catheter ablation, contact force has been demonstrated to be important for obtaining a permanent transmural lesion, which minimizes conduction recovery and associated clinical recurrences^11,12^. Improved outcomes may be due to patient immobility, improved accuracy of mapping, and catheter stability, allowing better contact between the catheter and the endocardium and optimizing lesion quality. In our study, only a small minority of cases were performed with contact sensing catheters as this technology only became available towards the end of the study period. It would however be useful to conduct further studies comparing average contact force between cases. Other studies underline the advantages of high-frequency jet ventilation to achieve better lesions during ablation^13,14^. Use of a remote magnetic navigation system for AF ablation reversed the improvement in outcome following GA^15^. There is also some evidence from a prospective non-randomized trial that AF patients prefer GA^16^.

We did not find any difference in procedure time between procedures; it is likely that the increased time taken to anaesthetise the patient and then wake them at the end of the procedure is similar to the time saved during the procedure itself in easier catheter manipulation. The actual ablation time was also not significantly different between procedure types.

There has been concern that atrio-oesophageal fistula may be more common when the procedure is performed under GA^17^; no instances occurred in our study in either group. No patients developed complications related to the GA; conversely 5 procedures under sedation were hindered by airway problems, patients becoming agitated or with uncontrolled pain.

In our study there was a significant decrease in the number of patients being listed for repeat procedures. This meant that even though conducting a procedure under GA is slightly more expensive than under sedation, the cost per patient after a maximum of two procedures is lower if all persistent AF ablations are conducted under GA.

The main limitations of this study lie in it being a retrospective analysis. It is therefore not possible to control for different operator technique and decisions based on individual patient factors. Additional randomized studies should be performed to confirm these results. For the cost-effectiveness analyses, the costs for GA provision in our centre have been used; however, costs may differ between centres and may need to be assessed locally. In some areas it would have been useful to have more robust data collection, for example on adverse events. Continuous monitoring was not employed to monitor for possible recurrence of paroxysmal AF. Furthermore, it would have been useful to gather Quality of Life information from patients undergoing AF ablation to better analyse the impact of ablation of the patient.

We have no funding declarations or conflicts of interest.

